# Altered hypothalamic DNA methylation and stress-induced hyperactivity in a novel model of early life stress

**DOI:** 10.1101/2020.04.09.033951

**Authors:** Eamon Fitzgerald, Matthew C Sinton, Sara Wernig-Zorc, Nicholas M Morton, Megan C Holmes, James P Boardman, Amanda J Drake

**Affiliations:** University/British Heart Foundation Centre for Cardiovascular Science, University of Edinburgh, The Queen’s Medical Research Institute, 47 Little France Crescent, Edinburgh EH16 4TJ, UK; Department of Biochemistry III, University of Regensburg, University of Regensburg, 93040 Regensburg, Germany; MRC Centre for Reproductive Health, University of Edinburgh, The Queen’s Medical Research Institute, 47 Little France Crescent, Edinburgh, EH16 4TJ, UK; Centre for Clinical Brain Sciences, University of Edinburgh, Chancellor’s Building, 49 Little France Crescent, Edinburgh EH16 4SB

## Abstract

Early life stress during childhood is associated with a number of psychiatric disorders that manifest across the life course. Preterm birth is a profound stressor, and an important cause of cognitive impairment, as well as neurodevelopmental and psychiatric disorders. However, the mechanisms that link events during the early neonatal period with later functional problems are poorly understood. We developed a novel mouse model of early life stress (modified maternal separation; MMS) with specific relevance to preterm birth (PTB) and hypothesised it would affect the hypothalamic transcriptome and DNA methylome and impact on behaviour in adulthood. MMS consisted of repeatedly stimulating pups for 1.5 hours/day, whilst separated from their mother, from postnatal day (P)4-6. 3’ RNA sequencing and DNA methylation immunoprecipitation (meDIP) sequencing was performed on the hypothalamus at P6. Behaviour was assessed with the elevated plus and open field mazes, and in-cage monitoring at 3-4 months of age. Although MMS was only associated with subtle changes in gene expression there were widespread alterations in DNA methylation. Notably, differentially methylated regions were enriched for synapse-associated loci. MMS also resulted in hyperactivity in the elevated plus and open field mazes, but in-cage monitoring revealed that this was not representative of habitual hyperactivity. In conclusion we describe a novel model of early life stress with relevance to PTB, with marked effects on DNA methylation in the hypothalamus and with stress-specific hyperactivity in young adulthood. We suggest that these results have implications for the understanding of early life stress mediated effects on brain development.

## Introduction

Preterm birth (PTB), defined as birth prior to 37 weeks of gestation, accounted for 10.6% of total births worldwide in 2014 [1]. The rate of PTB has increased annually since 2014 in the United States [2]. Although mortality and morbidity have decreased amongst infants born preterm and very low birthweight in many high income countries [3], PTB remains the largest cause of morbidity and mortality in the neonatal period [4]. One of the major long-term consequences of PTB is cognitive impairment in childhood [5], ranging from severe motor disorders to cognitive abnormalities [6], and PTB is closely associated with autism spectrum disorder (ASD) [7], schizophrenia [8], attention deficit hyperactivity disorder (ADHD) [9] and various psychiatric disorders in later life [10].

Early life stress has been associated with a number of chronic health problems, including psychiatric disorders [11]. Infants born preterm often have extended stays in the neonatal intensive care unit (NICU), and this is associated with a number of environmental stressors; such as exposure to exogenous stimuli (e.g. light and noise), invasive and/or painful procedures and sleep deprivation [12, 13], as well as various physiological stressors [14]. Studies designed to investigate the contribution of such stressors to the long-term effects of PTB are difficult, because of the presence of multiple confounders such as illness severity or length of the NICU stay. Nevertheless, studies which have attempted to correct for various confounders have shown that the number of painful procedures experienced associates with alterations in brain development [15–17] suggesting that some stressors may indeed have long term consequences. Moreover, environmental enrichment in the form of music can improve the functional brain architecture of the preterm brain [18], and improved nutrition through the use of breast milk feeding is associated with improved brain connectivity [19] although the precise mechanisms are unknown.

It has been proposed that early life stress may mediate some of its effects on neurodevelopmental outcome through alterations in epigenetic modifications including DNA methylation [20]; for example, the association between childhood abuse and post-traumatic stress disorder may be mediated through changes in DNA methylation [21]. This may also have relevance for preterm birth, indeed we have shown alterations in DNA methylation at key neurodevelopmental loci in buccal cells of infants born preterm, which were taken at term equivalent age [22]. A number of further studies have reported differences in DNA methylation between preterm and term infants at birth [23–26] although the extent to which these are persistent is unclear [25, 27–29]. To better understand the relative influence of potential stressors associated with PTB, and to dissect underlying mechanisms, including epigenetic dysregulation in the tissue of interest, a number of animal models have been used [30]. Various early life stress paradigms, including maternal separation and altered maternal care have been shown to associate with an array of different phenotypes, including effects on the hypothalamic-pituitary adrenal (HPA) axis and vulnerability to adult stress [30]. These studies have reported differences in DNA methylation at loci which may impact on HPA axis feedback [30], for example at the *arginine vasopressin* (*AVP*) locus in the hypothalamus [31].

In the most commonly used model of early life stress, maternal separation, pups are separated from the mother for 3 or more hours per day for 10 or more consecutive days. Considering the mother is the sole source of nutrition during this time, nutritional deficits could confound any findings in this model. Indeed, we have previously shown that extended periods of maternal separation in rats results in hypoglycaemia [32], which is independently associated with impaired neurodevelopment [33]. Further, traditional maternal separation models may not accurately recapitulate some of the common stressors associated with the NICU, including frequent handling and sleep deprivation. We therefore developed a novel model of early life stress, termed modified maternal separation (MMS), involving short periods of maternal separation in combination with manual manipulation, which we propose has relevance to infants on the neonatal intensive care unit. Since in mice, brain development at birth is roughly equivalent to that of a human at 24 weeks post-conception and matures to term equivalence by postnatal day (P)10 [34], MMS occurs at time-points with neurodevelopmental relevance to PTB in humans.

Given the central role of the hypothalamus as a mediator of the response to early life stress [31, 35], we hypothesised that MMS would result in perturbations of the hypothalamic transcriptome and DNA methylome and in altered behaviour in adulthood. We tested this using a combination of specific candidate gene expression analysis (for genes involved in glucocorticoid signalling and DNA methylation), 3’ mRNA sequencing and DNA methylation immunoprecipitation (meDIP) sequencing in the hypothalamus at P6. We then performed behavioural assessment using the elevated plus maze (EPM), open field (OF) and in-cage behavioural analysis of habitual activity at 3-4 months of age, in combination with candidate gene expression and DNA methylation analysis in adult mice. We found that MMS associates with profound changes in hypothalamic DNA methylation in the neonatal period and with a stress-specific hyperactive phenotype in adulthood.

## Methods

### Animals

All experiments were carried out under University of Edinburgh guidelines under the UK Home Office Animals (Scientific Procedures) Act 1986. Adult C57/Bl6 mice (Harlan, UK) had *ad libitum* access to chow (Special Diets Services, Essex, UK) and water (lights on 07:00-19:00, temperature 22°C). For mating, 2 females and 1 male were kept per cage. P0 was designated as the day of birth. Mice were weaned at P21 with littermates housed together. Neonatal experiments consisted of n=10/group, unless otherwise stated. Adult experiments were performed on n=11/group for biochemistry, elevated plus maze (EPM), open field (OF) and tail suspension and n=7 for in-cage behavioural analysis.

### Modified maternal separation (MMS) and behavioural testing

At P3, all female pups were killed. Four of the remaining males/litter were randomised to control or MMS. MMS was performed between 1330-1500 daily from P4-P6 in a separate room. MMS pups were placed on a heating pad adjacent to the home cage allowing communication through ultrasonic vocalisation. Pups were gently moved to the supine position whenever they returned to a prone position; this occurred continuously for 1.5 hours. Pups were then returned to the home cage. Pups were weighed daily; weights were normalised to starting weight at P4.

Immediately after MMS on P6, one cohort was killed by decapitation. Trunk blood was collected, and blood glucose measured immediately using the AccuCheck Performa Glucometer (Roche, UK). Whole blood was collected in EDTA coated tubes (Sarstedt, Germany) and plasma isolated and stored at −80°C. Whole brains were extracted and the hypothalamus dissected. Tissue was snap-frozen on dry ice and stored at −80°C or fixed in 4% paraformaldehyde.

A second cohort of mice was weaned at P21, with control and MMS littermates housed together. Behavioural testing (EPM, OF and tail suspension) was performed at P90-P100. The EPM was used as previously described [48]. Mice were placed in the centre zone facing the open arm and left to explore the maze freely for 5 minutes. The OF test was carried out as previously described [49], 24 hours after the EPM. Mice were placed in the centre of the OF and allowed to freely explore the environment for 5 minutes. For EPM and OF, mice were habituated to the novel holding area for at least one hour, the tester was hidden from view. Recording and analysis was performed automatically using the AnyMaze software (AnyMaze, Dublin). One hour after the OF, tail suspension testing was performed as previously described [50] for 6 minutes with the tester out of sight. Recordings were analysed for time immobile with the investigator blinded to group.

At P120-P130, in-cage behaviour was analysed using the TSE-systems PhenoMaster (TSE-Systems, Germany). Mice were single housed for 4 days to acclimatise and then introduced into a fresh home cage and allowed to acclimatise for 24 hours. Activity measurements were taken automatically every 15 minutes over two consecutive 24-hour periods and then averaged. Animals were killed by cervical dislocation; tissue was dissected, frozen on dry ice and stored at −80°C.

### Corticosterone ELISA

Blood samples were collected for corticosterone analysis by tail venesection at 7am and 7pm the day after the tail suspension test. Plasma corticosterone was analysed by ELISA (Enzo Life Sciences, Exeter).

### Immunohistochemistry

Brain tissue was fixed overnight in 4% paraformaldehyde, before transferring to 70% ethanol overnight and embedding in paraffin wax. 5μm sections were cut using a Leica 2M2125 RTS microtome (Leica, Milton Keynes) and three consecutive sections placed on each slide, dried overnight at 37°C and stored at room temperature. Primary antibodies (Supplementary Table 1) were applied overnight in a humid chamber at room temperature. Slides were washed in PBS, secondary antibody applied for 1.5 hours and washed in PBS before mounting media and coverslips were applied. Slides were imaged using the Zeiss AxioScan Z1 (Zeiss, Cambridge) system using a 10x objective. Files were exported with original metadata to a Tiff format before downstream analysis with Image J.

### DNA/RNA extractions

DNA/RNA were extracted from the same sample using the Qiagen All Prep DNA/RNA Mini Kit (Qiagen, Manchester).

### Reverse transcription and quantitative PCR

1µg of RNA was DNase treated with RQ1 RNase-free DNase (Promega, Hampshire). Reverse transcription was performed with the Applied Biosystems RT kit (Thermo Fisher Scientific, UK) in a G-Storm Thermocycler (Akribis Scientific Limited, Cheshire). qPCR primers were designed using the UPL assay design centre and cDNA samples analysed on a Roche LightCycler 480, normalised to the expression of the housekeeping gene *TBP*. Primers are listed in Supplementary Table 2.

### 3’ mRNA sequencing

For 3’mRNA sequencing, 6 samples were randomly selected per group. Sequencing was performed at the Wellcome Trust Clinical Research Facility (University of Edinburgh). Library preparation was done using the QuantSeq 3’ mRNA-Seq Library Prep Kit (Lexogen, Austria). The template was prepared using the Ion PI Hi-Q OT2 200 kit (Thermo Fisher Scientific, UK). Sequencing was performed using the Ion PI Chip Kit v3 (Thermo Fisher Scientific, UK), 12 samples per chip. The Ion Hi-Q Sequencing 200 Kit (Thermo Fisher Scientific, UK) and the Ion Proton platform were used for analysis. For data analysis, raw pH DAT files were converted to flow signals and aligned to the mm10 reference genome in an automated workflow (Torrent Suite v 5.2.0). Analysis was performed using Galaxy [36] and Degust software. Gene Ontology was carried as previously described [37, 38], using genes with a log fold change of greater than 1.5 [39, 40]. Transcription factor enrichment analysis was carried out using the oPOSSUM-3 software as previously described, with a Fisher score <7 and a Z-score <10 taken as significant enrichment [41]. Data are available through the Gene Expression Omnibus (GSE147375).

### DNA methylation immunoprecipitation and sequencing

For DNA methylation immunoprecipitation (meDIP)-sequencing, three samples/group consisting of three pooled biological replicates were sequenced as described previously, using the Ion Proton platform [42]. A mean read length of 133-144 base pairs and 24,288,817-34,030,252 reads per sample was achieved. Reads were aligned to the MM10 genome using Torrent Suite v5.2.0. Aligned reads were sorted using SAMtools, before calling peaks using MACS2 (v. 2.1.1) -f BAM --broad --broad-cutoff 0.05 -B -g hs, over corresponding inputs [43]. To detect differentially methylated regions (DMRs), we used Diffbind with DESeq2 and edgeR options [44]. DMRs were assigned to genes and other genomic features using the HOMER (v. 4.8) annotatePeaks tools [45]. For sequencing, data were normalised to a pooled input for each group and subtracted from an IgG control. For candidate meDIP analysis, the concentration of each sample was extrapolated from a standard curve of arbitrary concentrations and normalised to 10% input. Regions of interest were identified from the meDIP-sequencing dataset. Primers were designed using the NCBI primer-BLAST software (Supplementary Table 3). Data are available through the Gene Expression Omnibus (GSE146892).

### Statistical analysis

All statistical analyses were performed using IBM SPSS software version 24. An independent t-test was used for candidate comparisons between control and MMS groups at a single timepoint. A repeated measures ANOVA was used for the comparisons of control and MMS groups at multiple timepoints (i.e. AM and PM).

## Results

### MMS is not associated with alterations in weight gain, blood glucose or plasma corticosterone

There were no differences in early weight gain, P4-P6 (Figure 1A), blood glucose concentrations (Figure 1B) or plasma corticosterone concentrations (Figure 1C) between groups.

**Figure 1.**
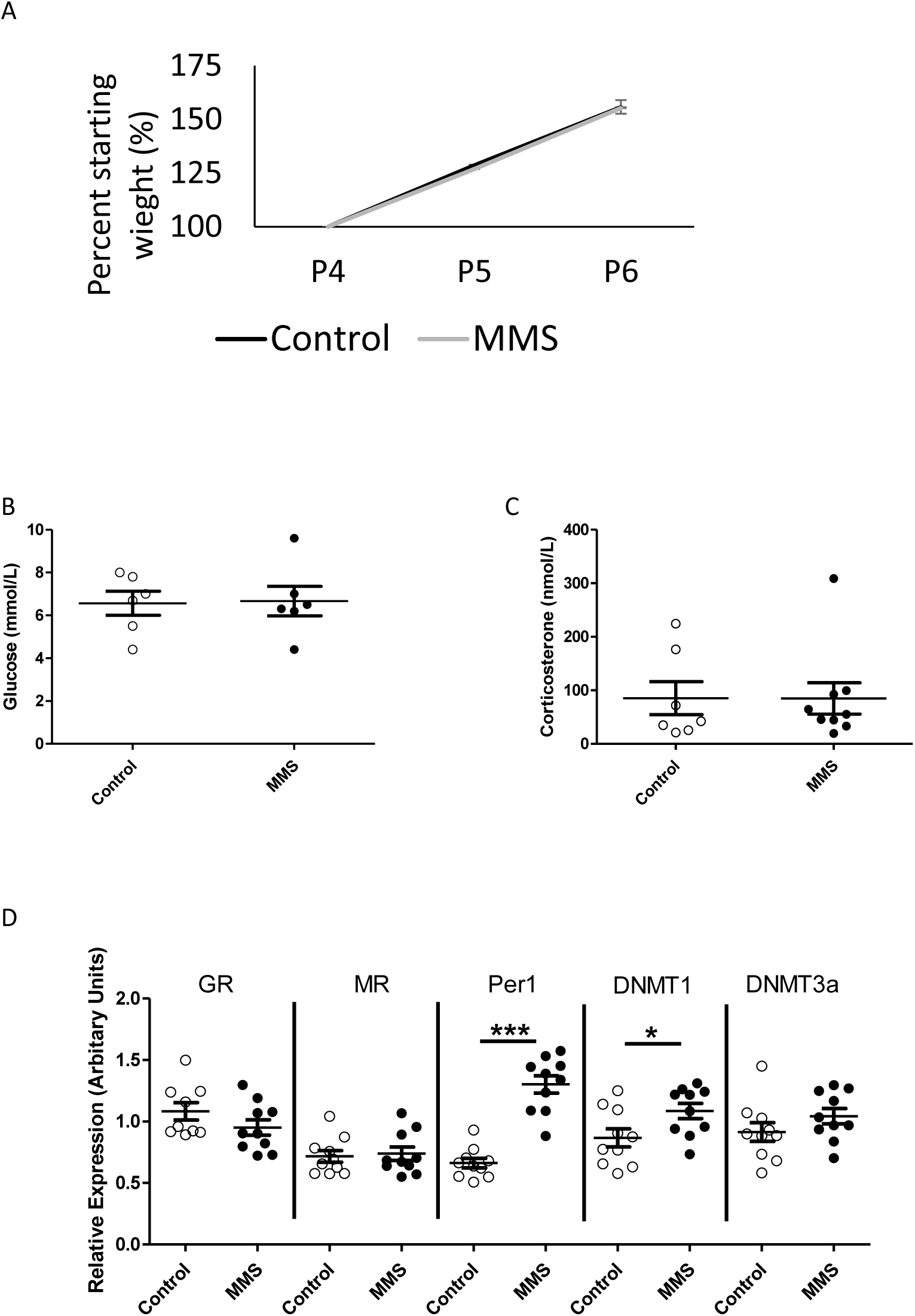
MMS is not associated with alterations in weight gain, blood glucose or plasma corticosterone, but is associated with changes in candidate gene expression in the hypothalamus. (A) There was no difference in weight gain between control (black solid line) and MMS (orange dashed line) groups during MMS (p=0.94 for area under the curve), n=30/group. There were no differences between control (clear circles) and MMS (black filled circles), in (B) blood glucose (p=0.91) (n=6/group) or (C) plasma corticosterone concentrations (p=0.29) immediately following MMS (n=7 and 10/group). (D) MMS was associated with increased mRNA expression of *Per1* (p=0.009) and *DNMT1* (p=0.03) but there were no differences in the expression of *GR* (p=0.053), *MR* (p=0.49) or *DNMT3a* (p=0.09), n=10/group. All statistical comparisons were made by independent t-test. Error bars indicate standard error of the mean.

### MMS is associated with early changes in hypothalamic gene expression

Analysis of hypothalamic candidate gene expression for genes involved in glucocorticoid signalling and DNA methylation revealed increased expression of the glucocorticoid responsive gene *Period 1* (*Per1*), which is associated with circadian rhythms, and *DNA methyltransferase 1* (*DNMT1*) (Figure 1D). There were no changes in the expression of the *glucocorticoid receptor* (*GR*), *mineralocorticoid receptor* (*MR*) or *DNA methyltransferase 3a* (*DNMT3a*). Considering the level of cellular heterogeneity within the hypothalamus and their responsiveness to stress in adult animals [51–53], we used immunohistochemistry to assess the distribution of ionized calcium binding adaptor molecule 1 (Iba1, a marker of myeloid cells), Olig2 (a marker of cells of the oligodendrocyte lineage) and glial fibrillary acidic protein (GFAP, which is increased in reactive astrocytes) in the hypothalamus. There were no differences between control and MMS groups (Supplementary Figure 1).

We next used 3’ mRNA sequencing to look at the hypothalamic transcriptome in an unbiased manner (n=6/group). Analysis using the Voom/limma method identified only one gene (*D630033O11Rik*, labelled in Figure 2A) as differentially expressed (FDR<0.05). Multi-dimension scaling (MDS) showed no clustering of groups (Supplementary Figure 2A). Next, we took those genes with a log fold change (FC) of >1.5 [39, 40] (Figure 2A: blue=downregulated, red=upregulated) and used Gene Ontology analysis to indicate processes which may be implicated in subtle changes in gene expression (Figure 2B). We found enrichment for terms associated with “DNA binding RNA polymerase specific” and “Ligand-gated cation channel activity” within the molecular function category. Genes identified in these enrichment terms with the greatest fold change were *NR4A3* and *FOS*; upregulation of both of these genes was validated by qPCR (Figure 2C). Transcription factor binding enrichment analysis of genes with logFC>1.5 using the oPOSSUM-3 software [41], showed enrichment in motifs associated with several factors with key neurodevelopmental functions (Supplemental Figure 2B) including Nuclear Factor kappa B (NFkB) [54], Hypoxia Inducible Factor (HIF) 1α [55] and Krüppel-like factor 4 (KLF4) [56, 57].

**Figure 2.**
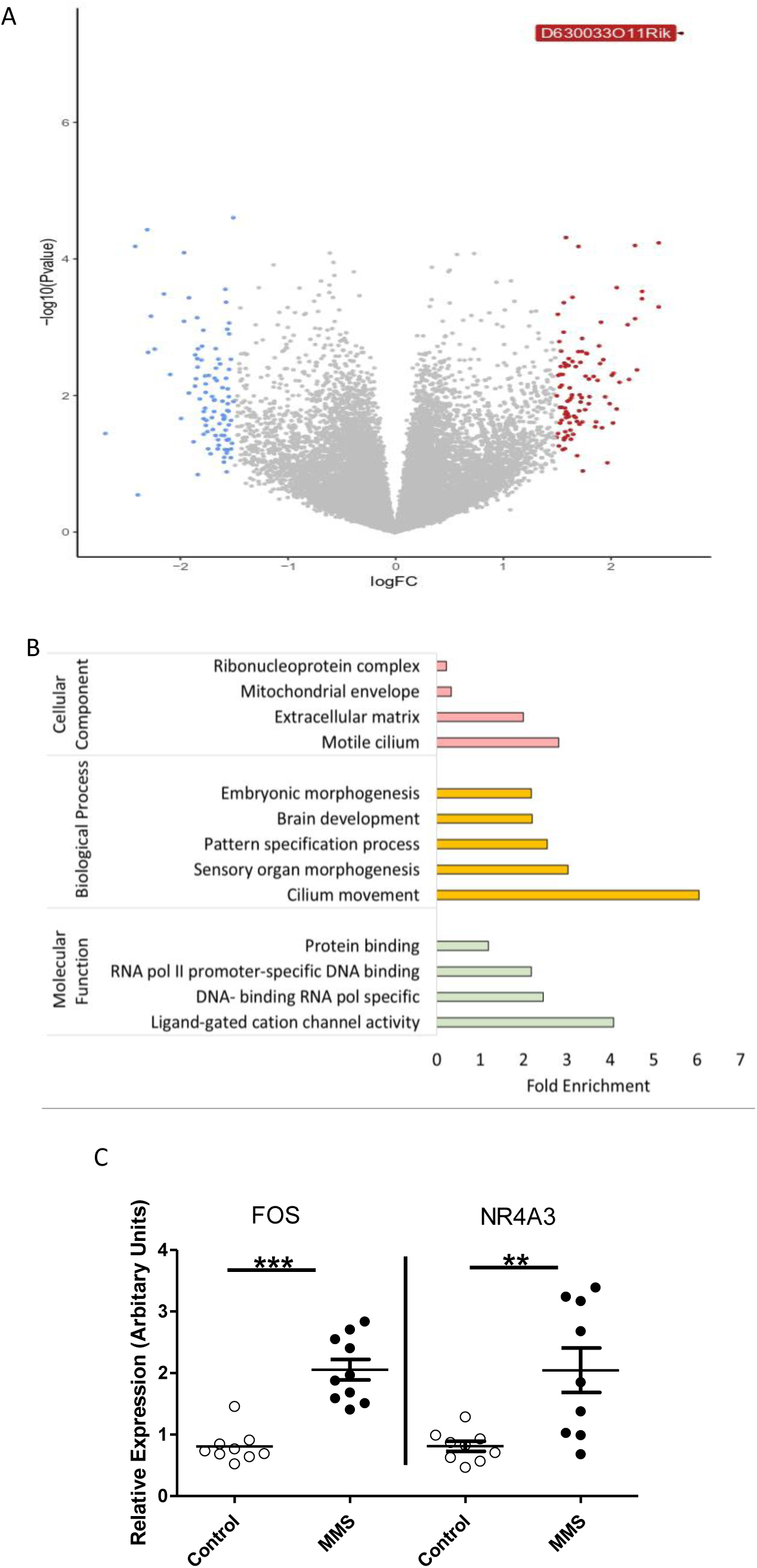
3’ mRNA sequencing of the hypothalamic transcriptome reveals only subtle changes associated with MMS. 6 samples were sequenced per group. (A) Volcano plot showing differential gene expression between groups, genes with a logFC greater than 1.5 are coloured blue (decrease) or red (increase). Only 1 gene (*D630033O11Rik*) had an FDR>0.05. (B) Gene Ontology analysis of genes with a logFC >1.5 (i.e. those coloured blue or red in A) for cellular component (pink), biological process (yellow) and molecular function (green). (C) Gene expression patterns, from control (clear circles) and MMS (black filled circles), identified from 3’ mRNA sequencing and Gene Ontology were validated using qPCR. FOS was enriched in the Gene Ontology term “Ligand-gated cation channel activity” and NR4A3 was identified from the enriched term “DNA-binding transcription activator activity, RNA polymerase II-specific”, in the biological process category. Both FOS (p>0.001) and NR4A3 (p=0.004) were significantly increased in expression following MMS. (C)

### MMS associates with early alterations in hypothalamic DNA methylation

Since expression of the DNA methylase *DNMT1* was increased following MMS, we postulated that DNA methylation might be affected. In total, nearly 13,000 DMRs were identified. Principal component analysis showed a distinct effect of MMS on the DNA methylome (Figure 3A). All DMRs are represented in the heatmap depicted in Figure 3B, clustered by Euclidian distance and within group samples were highly correlated (Figure 3C). Next, we identified DMRs associated with protein coding regions (Figure 3D). Gene Ontology analysis revealed enrichment for terms associated with various synaptic elements (Figure 3E). Differential methylation analysis was primarily performed using DESeq2, but there was a substantial overlap with DMRs identified using an alternative method (edgeR) (Supplementary Figure 3A).

**Figure 3.**
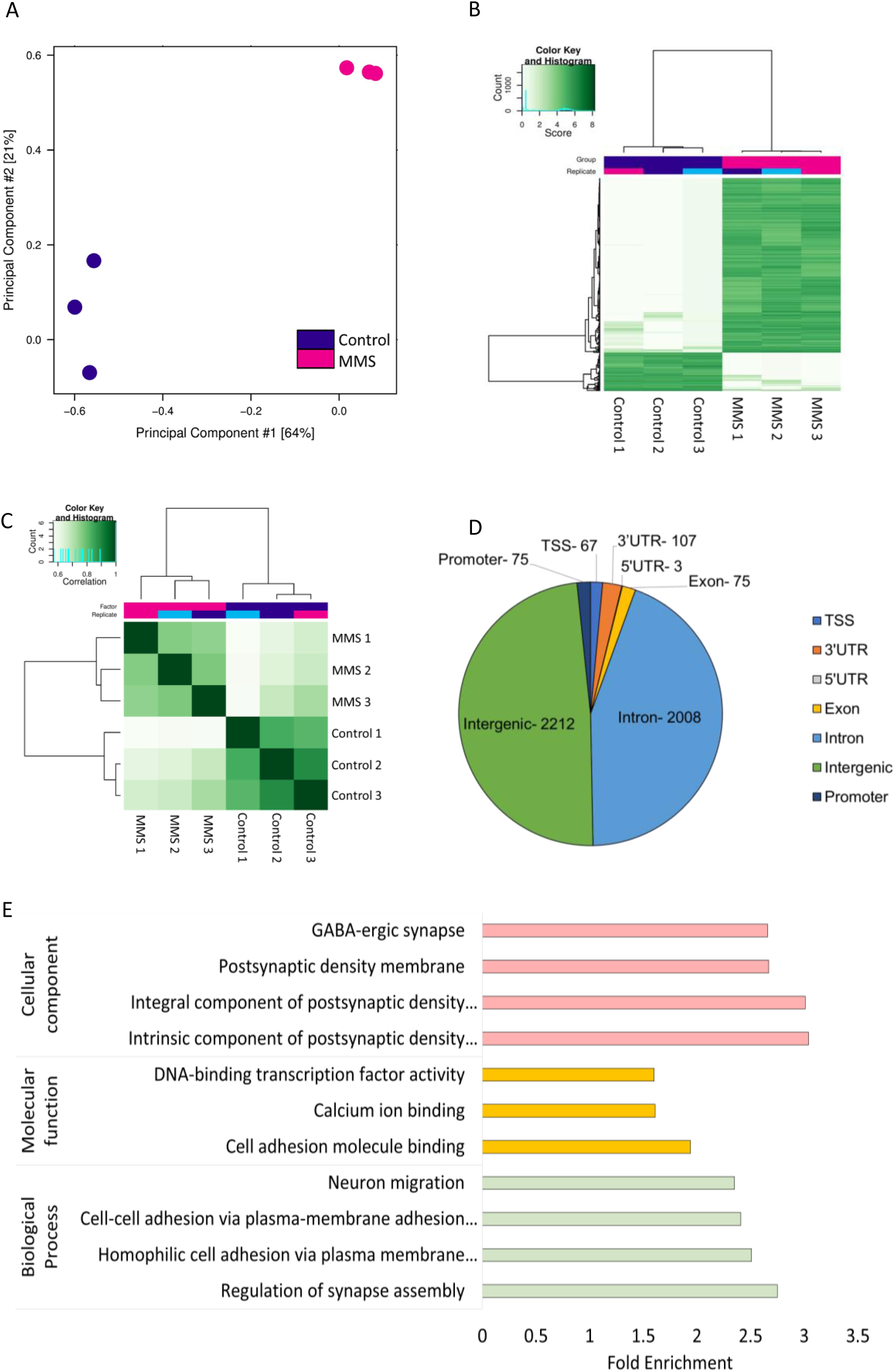
Widespread alterations in DNA methylation within the hypothalamus following MMS using meDIP sequencing. 3 samples were sequenced per group. (A) Principal component analysis of differentially methylated regions shows distinct clustering of control (blue) and MMS (pink) groups. Principal components 1 and 2 accounted for 64% and 21% of variance in the dataset respectively. (B) Heatmap of all DMRs throughout the genome with an FDR<0.05, which are clustered by Euclidian distance. (C) Correlation map of control and MMS samples following meDIP sequencing clustered by Euclidian distance. (D) Catalogue of the DMRs associated with protein coding regions. Numbers indicate numbers of DMRs associated with that region. (E) Gene Ontology analysis of differentially methylated sites (FDR<0.05) associated with protein coding regions. Terms for cellular component (pink), molecular function (yellow) and biological processes (green) are shown. Notably, there is enrichment of synapse associated terms under cellular component and biological processes.

There was no correlation between DMRs associated with protein coding regions and transcript expression (Supplementary Figure 3B). In particular, although DNA methylation in promoter regions is classically associated with gene expression [58], fewer than 20 promoter DMRs were identified, and these were not associated with transcript expression.

### MMS is associated with hyperactivity in the elevated plus maze (EPM) and open field maze (OF) but not during habitual in-cage movement in adulthood

Representative track plots for control and MMS animals in the EPM are shown in Figures 4A and 4B. Mice exposed to MMS travelled further in the EPM (Figure 4C) but did not show any altered preference for the open arms, closed arms or centre zone (Figure 4D). The MMS group also had a higher overall average speed and a higher total mobile time (Supplementary Figure 4A and B). The MMS group also travelled further in the closed arm, with no difference in the duration of visit (Supplementary Figure 4C and D), while there was no difference between groups in distance travelled in the open arms or in the duration of each visit (Supplementary Figure 4E and F). Representative track plots for the control and MMS groups in the OF are shown in Figure 4E and 4F. MMS mice travelled further during the test (Figure 4G) but did not show an altered preference for the inner or outer zone (Figure 4H). MMS mice also moved at a higher speed and spent more time mobile (Supplementary Figure 5A and B). The MMS group travelled further in both the inner and outer zones of the OF, with a shorter length of visit to the outer zone but no difference in length of visit to the inner zone (Supplementary Figure 5C-F). There were no differences between groups in the tail suspension test (Supplementary Figure 5G).

**Figure 4.**
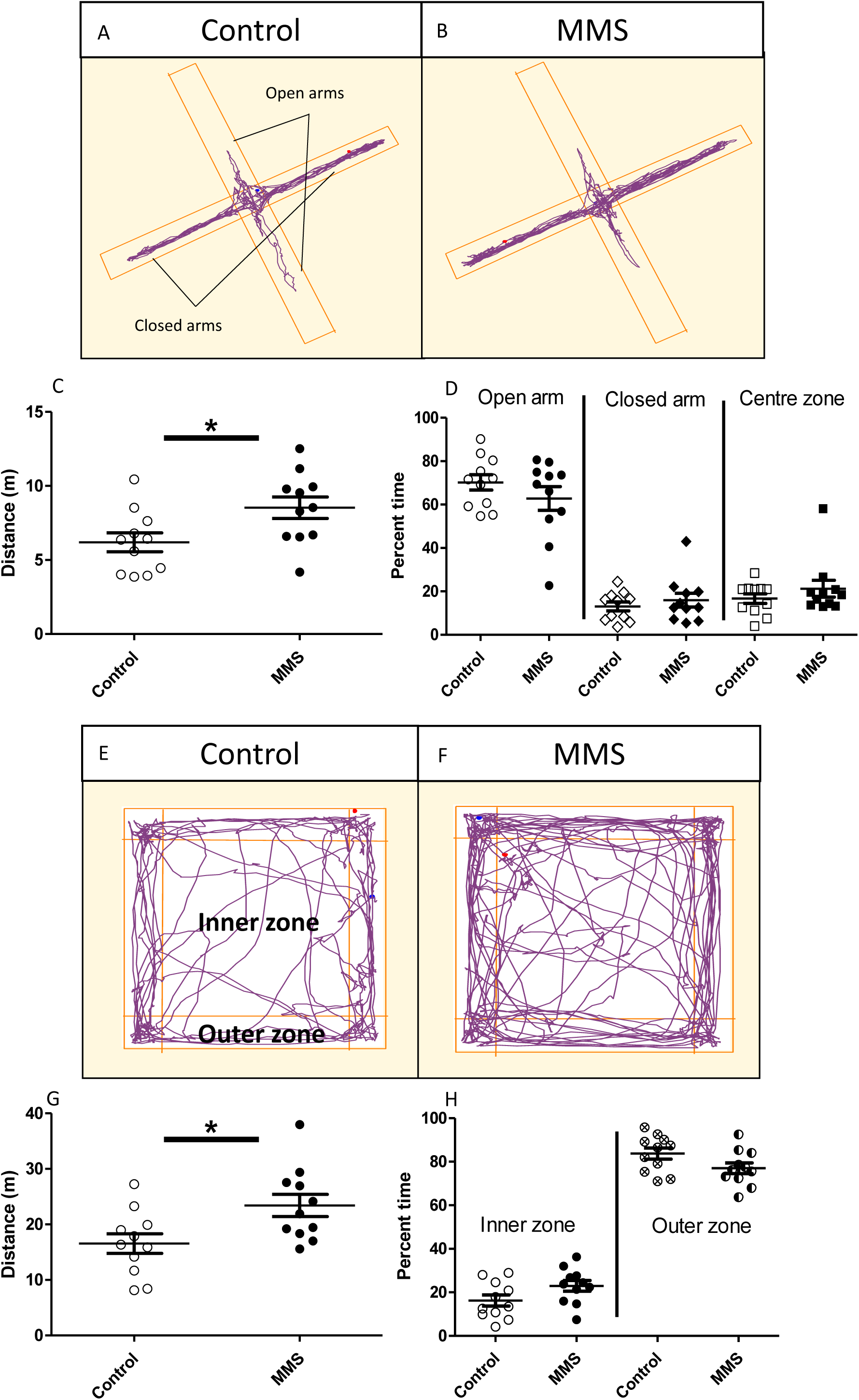
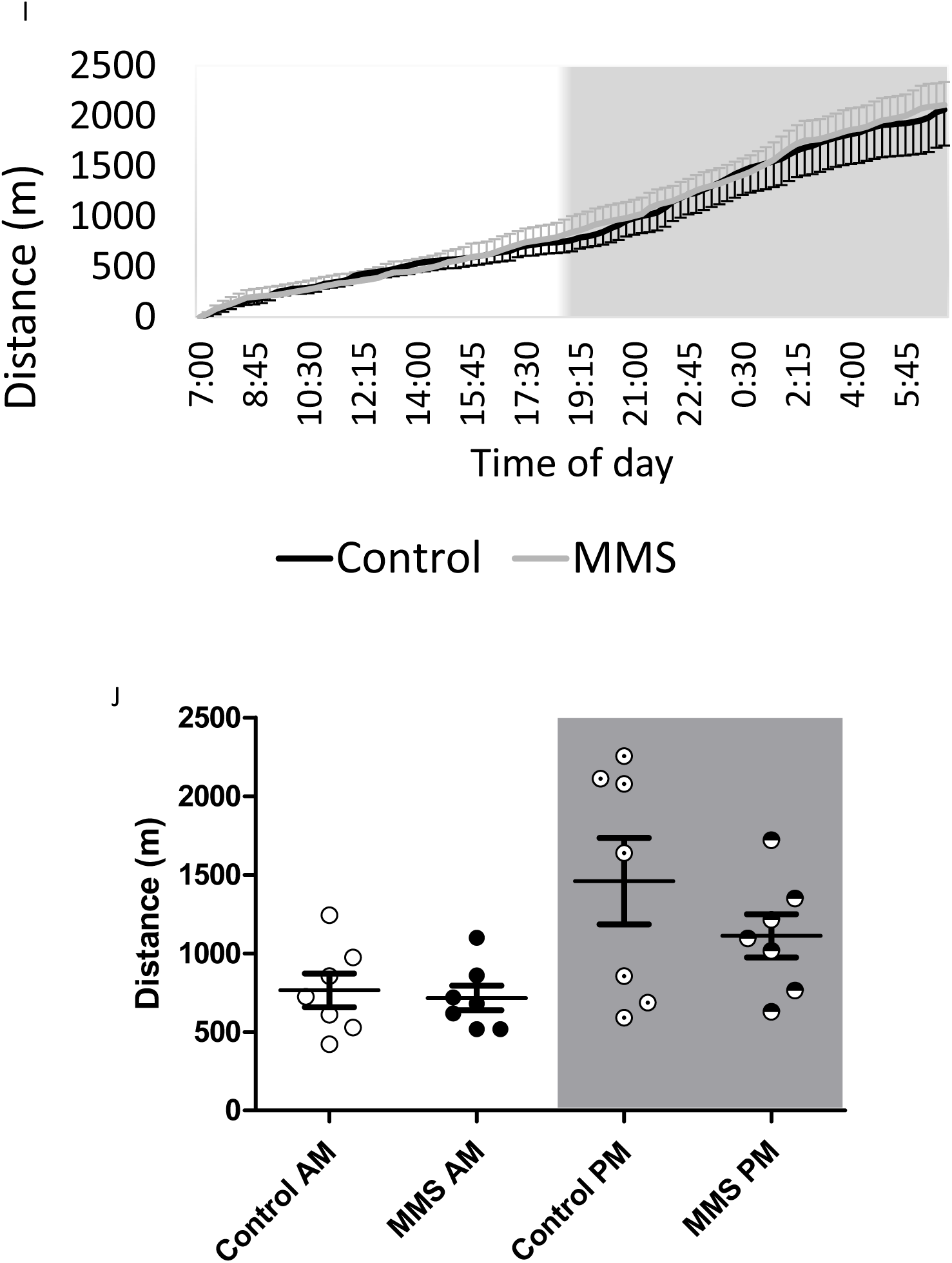
MMS is associated with hyperactivity in the elevated plus (EPM) and open field (OF) maze but not during habitual in cage movement. (A and B) Representative track plots for control and MMS groups in the EPM, with the open and closed arms of the maze indicated in A. (C) The MMS group travelled further during the EPM (n=11/group) (p=0.02), but there were no differences in the time spent in the open, closed or centre areas (D). (E and F) Representative track plots for control and MMS groups in the OF with the inner and outer zones indicated in E. (G) The MMS group travelled further in the OF (n=11/group) (p=0.02) and spent similar amounts of time in the inner and outer zones (H). (I and J) In-cage behavioral monitoring using the TSE PhenoMaster system tracked movement over a 24-hour period (two consecutive 24-hour periods were averaged for each animal). (I) Cumulative distance travelled with respect to time of day, the light aspect of the graph indicates when the lights were on and the dark aspect indicates when the lights were off. (J) Total movement calculated for the light and dark phase (indicated by AM and PM, as well as the light and dark aspects) revealed no difference between the groups (p=0.36) (n=7/group) and no interaction with time (p=0.35). As expected, there was, a significant effect of time such that animals were active during the dark phase (p=0.001). Independent t-tests were used for C, D, G and H, while a two-way ANOVA with repeated measures was used to assess statistical associations in J. Error bars indicate standard error of the mean.

The EPM and OF constitute novel stressful environments. As such our next question was whether this hyperactivity was representative of higher habitual levels of activity or if it was specifically related to the novel stressful environment. To evaluate this, we used the TSE PhenoMaster in-cage monitoring system. There were no differences in habitual movement between groups either in terms of cumulative movement (Figure 4I), movement specifically during light and dark phases (Figure 4J), or in movement associated with grooming behaviours (Supplementary Figures 6A and B). Detailed analysis of other habitual and circadian behaviour patterns showed no differences between groups in food intake, calorie expenditure and fuel usage (Supplementary Figures 6C-E). Finally, there were no differences in plasma corticosterone, total body weight, lean body mass or fat mass (Supplementary Figures 7A-D).

### MMS does not associate with persistent changes in candidate gene expression or DNA methylation in the hypothalamus in adulthood

Considering the stress induced hyperactivity induced following MMS, we next evaluated the expression of genes associated with stress signalling in the adult hypothalamus. We identified no changes in the expression of *GR, MR, Per1* or *FKBP5* (Figure 5A). Further, there were no changes in the expression of stress-associated genes elsewhere in the hypothalamic-pituitary-adrenal axis including in the adrenal or pituitary gland (Supplementary Figure 8A and B). We did however find changes in expression of the *GR, FKBP5* and *Per1* in the hippocampus (Supplementary Figure 8C).

**Figure 5.**
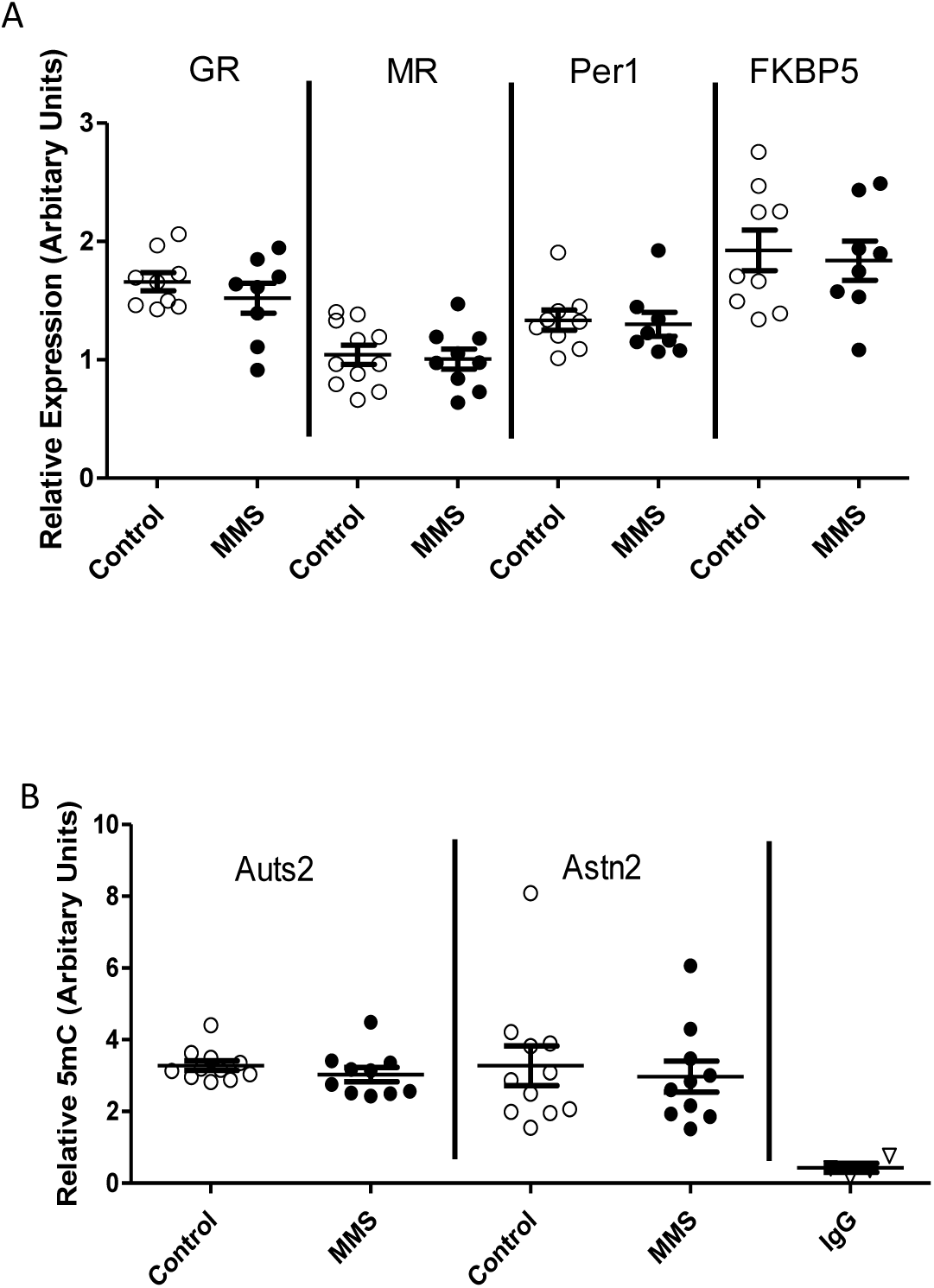
No persistent changes in candidate gene expression or DNA methylation in the hypothalamus at 4 months following MMS. (A) There were no differences in expression of the *GR* (p=0.35), *MR* (p=0.76), *Per1* (p=0.80) or *FKBP5* (p=0.72) in the hypothalamus between control (clear circles) and MMS (black filled circles) mice at 4 months of age. (B) There were also no differences in DNA methylation at candidate loci (*Auts2*: p=0.31, *Astn2*: p=0.68) in the hypothalamus between control (clear circles) and MMS (black filled circles). All statistical comparisons were made using an independent t-test. Error bars indicate standard error of the mean, with an n=7-11 for all groups.

Finally, we investigated DNA methylation within the hypothalamus at two loci which have been implicated in neurodevelopmental disorders, *Auts2* and *Astn2* [59, 60] and which were differentially methylated at P6 (Figure 3). We found no evidence for differential methylation at these loci in the hypothalamus at 4 months of age (Figure 5B).

## Discussion

We present a novel model of early life stress with relevance to PTB. The repetitive manipulation is intended to better model the manipulation and sleep deprivation associated with stays in the NICU [12]. To the best of our knowledge this is the first study to show a dichotomy between changes in the transcriptome and DNA methylome in the hypothalamus following early life stress, using unbiased sequencing-based approaches from the same samples.

Although animal models can be useful in furthering our understanding of the long-term effects of early life stress and in facilitating the dissection of underlying mechanisms, large differences in experimental design and differing behavioural outcomes have made interpretation difficult [61]. Previous studies using traditional maternal separation paradigms have described conflicting effects on behaviour [62, 63]. MMS is far shorter than all of these paradigms and indeed, the lack of effects on weight gain support that it is a relatively mild stressor and that maternal care is maintained. We also speculate that the active manipulation component of MMS may lead to more consistent experience of the stress between pups and thereby reduce heterogeneity in adult behavioural outcomes. Further, this model avoids hypoglycaemia as an additional confounder, which is important since hypoglycaemia is independently associated with atypical neurodevelopment in preterm infants [33].

We believe this is the first study to characterise in detail the habitual and circadian behaviours of mice exposed to early life stress. The behaviour differences observed in the EPM and OF are not mirrored by increased movement in the home-cage or any other alterations in in-cage circadian activities suggesting that the experience of a novel environment is specifically responsible for the hyperactivity during these tests. Preterm infants are at an increased risk of developing ADHD [9, 64], a pervasive neurodevelopmental disorder involving inattention and hyperactivity/impulsivity. Although the precise mechanisms linking preterm birth with ADHD are unknown, a number of hypotheses have been advanced, including early life environmental exposures, HPA axis dysregulation and perinatal inflammation [9]. Whether the ADHD phenotype in preterm infants is responsive to novel environments is not known, but abnormal responses to stress are certainly present in a number of other psychiatric and neurodevelopmental disorders including anxiety disorders [65], ASD [66], schizophrenia [67], bipolar disorder [68] and are well-described as a consequence of early life stress in humans [69].

Many psychiatric disorders are multifactorial and influenced by both genetic predisposition and environmental factors [70–73]. Adverse early life events have been associated with an increased risk of mood and anxiety disorders [74]; for example, genetic variants associated with schizophrenia interact with early life stress to affect risk [75]. Moreover, childhood stress increases the risk for later psychiatric disorders [11]: temporary childhood neglect is associated with an increased incidence of ADHD [76], and child abuse associates with suicidal ideation and various psychiatric disorders [77–79]. Moreover, structural and function networks which include the hippocampus and amygdala are altered following PTB [80–82]. Our data suggest that perinatal stress may contribute towards the atypical brain development associated with PTB, so that studies aimed at dissecting the specific consequences of early life stress on the preterm brain may increase our understanding of the pathogenesis of neurodevelopmental disorders in these individuals.

Glucocorticoids are a primary mediator of the physiological response to stress, but MMS did not result in increased plasma corticosterone concentrations at P6. The classic mechanism of glucocorticoid action involves transcriptional alterations following GR binding, and an initial candidate approach to assess gene expression revealed increased expression of the glucocorticoid sensitive gene and circadian regulator, *Per1*. However, transcriptome-wide analysis revealed minimal changes in gene expression, and transcription factor binding enrichment analysis of genes with logFC >1.5 did not show enrichment for glucocorticoid binding elements. This suggests a limited role for traditional glucocorticoid mediated transcriptional changes immediately following MMS, in line with the well-characterised stress hyporesponsive period present in neonatal rodents [83]. We were only able to measure basal and peak corticosterone in the home cage in adulthood and although we observed the expected diurnal variation there were no differences between groups. This is perhaps not surprising given the lack of differences in any of the measured in-cage parameters and supports the suggestion that the observed behaviours are induced by exposure to a novel environment but are absent at rest.

DNA methylation is highly dynamic during human brain development [84] and perturbations in DNA methylation are associated with several neurodevelopmental disorders, including ASD [85, 86], schizophrenia [87] and ADHD [88], and have also been documented following PTB [22]. We found substantial changes in DNA methylation in the hypothalamus immediately after MMS, and although this was not associated with widespread differences in gene expression, we did find a transcriptional signature of increased neuronal activation and an enrichment of DMRs in synapse-associated genes. In the adult brain neuronal activity is a potent modifier of DNA methylation [89], and DNA methylation has crucial roles in regulating synapse associated genes [90]. Thus, it is possible that increased MMS-induced neuronal signalling may be one mechanism driving the early DNA methylation changes. Analysis of DNA methylation at candidate loci suggests that these alterations may not persist into adulthood. Nevertheless, considering the dynamic nature of DNA methylation during brain development [84] and the important role of the hypothalamus in HPA axis regulation [35], it is possible that these early changes could be involved in programming the stress-specific hyperactivity in adulthood.

In conclusion, we have developed a novel paradigm of early life stress which models stress associated with the NICU. Using this model, we demonstrate substantial differential methylation in the hypothalamus in the perinatal period and stress induced hyperactivity at 3-4 months of age. We suggest that MMS is a useful model for the study of stress-associated alterations in brain development following PTB and provide novel insights into the progression and consequences of brain development following early life stress.

## Acknowledgments

This work was funded by a PhD studentship (to EF) from Medical Research Scotland (PhD-878-2015), in collaboration with Aquila BioMedical, Edinburgh, UK. MCS was supported by a British Heart Foundation PhD studentship (FS/16/54/32730) and by the British Heart Foundation Centre of Research Excellence. NMM was supported by a Wellcome Trust New Investigator Award (100981/Z/13/Z). MCH was supported by WT project grant (WT079009J. PB was supported by the MRC Centre for Reproductive Health Centre Grant (MRC G1002033). AJD was supported by the British Heart Foundation Centre of Research Excellence, University of Edinburgh. All sequencing was carried out at the Wellcome Trust Clinical Research Facility at the Western General Hospital, University of Edinburgh.

