## Supplementary tables and figures for "Altered hypothalamic DNA methylation and stress-induced hyperactivity in a novel model of early life stress"

### Supplemental table legends

**Supplemental Table 1**

List of antibodies used in this manuscript

**Supplemental Table 2**

List of qPCR primers and associated UPL probe used in this manuscript

**Supplemental Table 3**

List of primers for candidate meDIP investigation

### Supplemental figure legends

**Supplemental Figure 1. There were no differences in GFAP+, Iba1+ or Olig2+ cell proportions in the hypothalamus following at P6 following MMS**. In A and B, representative images of GFAP+ staining from the hypothalamus are shown. In C, GFAP+ cells are quantified as percent coverage in control (clear circles) and MMS (black filled circles) groups. In D and E, representative images from Iba1 (green) and DAPI (blue) stained hypothalamic sections are shown, which are quantified in F as cells/$mm^2$ for control (clear circles) and MMS (black filled circles) groups. In G and H, representative images of Olig2 (green) and DAPI (blue) stained hypothalamic sections are shown, which are quantified in I as Olig2+ cells/$mm^2$ for control (clear circles) and MMS (black filled circles) groups. Scale bar indicates 100µm. All statistical comparisons were made with an independent t-test. Error bars indicate standard error of the mean, n=7-9/group.

**Supplemental Figure 2. Transcription factor binding enrichment analysis and 3' mRNA seq validation indicate NFkB activation and neuronal activity following MMS.** (A) MDS plot for all sequenced samples (control-blue and MMS-orange). Dimension 1 accounted for 50% of the total variance in the dataset, while dimension 2 accounted for 15%. (B) Transcription factor binding enrichment analysis using the oPOSSUM software, identifies transcription factors with binding sites overrepresented among genes which have a logFC >1.5. Listed are the Fisher (clear bars) and Z scores (black bars) for each transcription factor. Statistical comparisons were done using an independent t-test. Error bars indicate standard error of the mean, n=10/group.

**Supplemental Figure 3. meDIP sequencing.** (A) Venn diagram of 2 methods of analysis for differential DNA methylation shows significant overlap between DESeq2 and edgeR. There were 493 and 7047 unique DMRs associated with the edgeR and DESeq2 methods respectively, with 12456 DMRss identified in both methods. (B) There was no correlation between DMRs (y-axis) and expression of corresponding genes (logFC >0.5; x-axis). Pearson correlation coefficient 0.07; p=0.36**.**

**Supplemental Figure 4. Movement in EPM.** (A) The MMS group (black circles) moved faster than the control group (clear circles) (p=0.02). (B) The MMS group spent more time mobile (p=0.02), total test duration 300 s. (C) The MMS group travelled further in the closed arm (p=0.048), but (D) there was no difference in the duration of each visit to the closed arm (p=0.27). There were no differences in the distance travelled in the open arms of the maze (E, p=0.38) or the average duration of visit to the open arms (F, p=0.90). All comparisons were made using an independent t-test. Error bars indicate standard error of the mean, n=11/ group.

**Supplemental Figure 5. OF and tail suspension tests.** (A) Animals in the MMS group (black circles) moved with a higher speed throughout the testing period when compared to controls (clear circles) (p=0.02). (B) The MMS group spent more time mobile (p=0.02), total test time 300s. (C) The MMS group travelled further in the outer zone (p=0.04). (D) The average visit to the outer zone was shorter in the MMS group (p=0.048). (E) The MMS group also travelled further in the inner zone (p=0.02). (F) There was no difference between groups with respect to duration of visit to the inner zone (p=0.48). (G) In the tail suspension test, there was no difference in total immobile time between groups (p=0.32). Total length of testing 360s. All comparisons were made using independent t-tests. Error bars indicate standard error of the mean, n=11/ group.

**Supplemental Figure 6. In cage analysis reveals no statistical differences in in-cage activities, feeding behaviour and calorie expenditure following MMS.** There were no differences in (A) x-axis or (B) y-axis related grooming behavior as quantified by laser beam breaks in the specified plane (p=0.96 and 0.73 respectively). (C) There was no difference in food intake quantified as total feeding during lights on (control AM and MMS AM) and lights off (control PM and MMS PM). There was a significant effect of time (p>0.001), but no effect of MMS (p=0.75) or interaction between time and MMS (p=0.15). (D) There was no difference in calorie expenditure normalised for lean body mass (p=0.99) or in fuel usage as indicated by respiratory exchange ratio (E; control (black line) and MMS (grey line)) over 24 hours (error bars indicate standard error of the mean). N=7/ group.

**Supplemental Figure 7. No difference in plasma corticosterone, overall weight, fat or lean body mass between control and MMS groups.** (A) There were no differences in plasma corticosterone concentrations (p=0.51). Data were analysed using a 2-way ANOVA, n=8-10/group. Light aspects of the graph indicate plasma corticosterone at 7am, with the dark aspect of the graph representing plasma corticosterone measured at 7pm. (B) Total body weight did not differ between control (clear circles) and MMS (black filled circles) groups (p=0.14). Analysis of body composition by TD-NMR revealed no difference in (C) lean mass (D) or total fat mass between control (clear circles) and MMS (black filled circles) groups. Data were analysed using an independent t-test; n=7/group. Error bars indicate standard error of the mean.

**Supplemental Figure 8. No changes in candidate gene expression in the adrenal or pituitary glands but there are changes in gene expression in the hippocampus at 4 months of age, following MMS.** (A) In the adrenal gland, there were no changes in gene expression for *Cyp11b1* (p=0.35), *MC2R* (p=0.72) or *stAr* (p=0.80) between the control (clear circles) and MMS (black filled circles) groups. (B) There were no changes in candidate gene expression in the pituitary gland for the *GR* (p=0.81), the *MR* (p=0.13), *FKBP5* (p=0.92), *Per1* (p=0.09) or *POMC* (p=0.78) between control (clear circles) and MMS (black filled circles) groups. (C) In the hippocampus, there was increased expression of *GR* (p=0.003), *FKBP5* (p=0.003) and *Per1* (p=0.004), but no change in expression of the *MR* (p=0.37) or *HSD11b1* (p=0.08). All candidate gene expression was normalized to the expression of *TBP*. All statistical comparisons were made using independent t-tests; n=11/group. Error bars indicate standard error of the mean.

Supplementary Table 1

| Antibody | Catalogue number | Dilution |
|---|---|---|
| IBA1 | 13481357 | 1 in 750 |
| GFAP | ab7260 | 1 in 2000 |
| Olig2 | NBP1-28667 | 1 in 1000 |
| DAPI | D9542 | 1 in 1000 |
| Anti-rabbit Alexa 488 | A-11034 | 1 in 750 |

Supplementary Table 2

| Gene | Forward primer sequence (5'→3') | Reverse primer sequence (5'→3') | Probe |
|---|---|---|---|
| TBP | gggagaatcatggaccagaa | gatgggaattccaggagtca | 97 |
| FKBP5 | tctgacaggccgtattccat | tcgtgaaagagaagggaactg | 20 |
| DNMT1 | gctaccagtgcacctttggt | atgatggccctccttcgt | 1 |
| DNMT3a | ggtgcactgaaatggaaagg | gaagaggtggcggatgac | 13 |
| Per1 | accactgagagcagcaagagt | ctcaggaggctgtaggcaat | 3 |
| GR | cagtgttttctaatggatattcaagc | ggagcacaccaggcagag | 10 |
| MR | gccggcatgaacttagga | ccctcttctgggctctgg | 5 |
| Fos | gggacagcctttcctactacc | agatctgcgcaaaagtcctg | 67 |
| Nr4a3 | tgcctgtcagcactgagtatg | tgcttgtgatcttggtgcat | 25 |
| CRH | gaggcatcctgagagaagtcc | tgttaggggcgctctcttc | 34 |
| POMC | gcttgcaaactcgacctctc | ttttcagtcaggggctgttc | 72 |
| CRHR1 | cttctccttctggggctga | aggtgccaatgaggtccac | 10 |
| HSD11b1 | tctacaaatgaagagttcagaccag | gccccagtgacaatcacttt | 1 |
| Cyp11b1 | agctcagacttggtgcttcag | gcccatggaatacagattcac | 3 |
| stAr | aaggctggaagaaggaaagc | ccacatctggcaccatctta | 2 |
| MC2R | caccacaatcctctaccctca | ggtgctgagaacttttccaataa | 55 |

Supplementary Table 3

| Associated gene | Forward primer sequence | Reverse primer sequence |
|---|---|---|
| Tex19.2 intragenic- Positive control | gggagatatgtaaatgagctgg | catccttacctccctgactgag |
| GAPDH promoter- Negative control | ccactccccttcccagtttc | cctataaatacggactgcagc |
| Auts2 | gtgctggggtcagaagacaa | tggcttgagttgagtgctgt |
| Astn2 | actcctcctttgggagggaa | cagtggtcagagtcagagagac |

Supplementary Figure 1

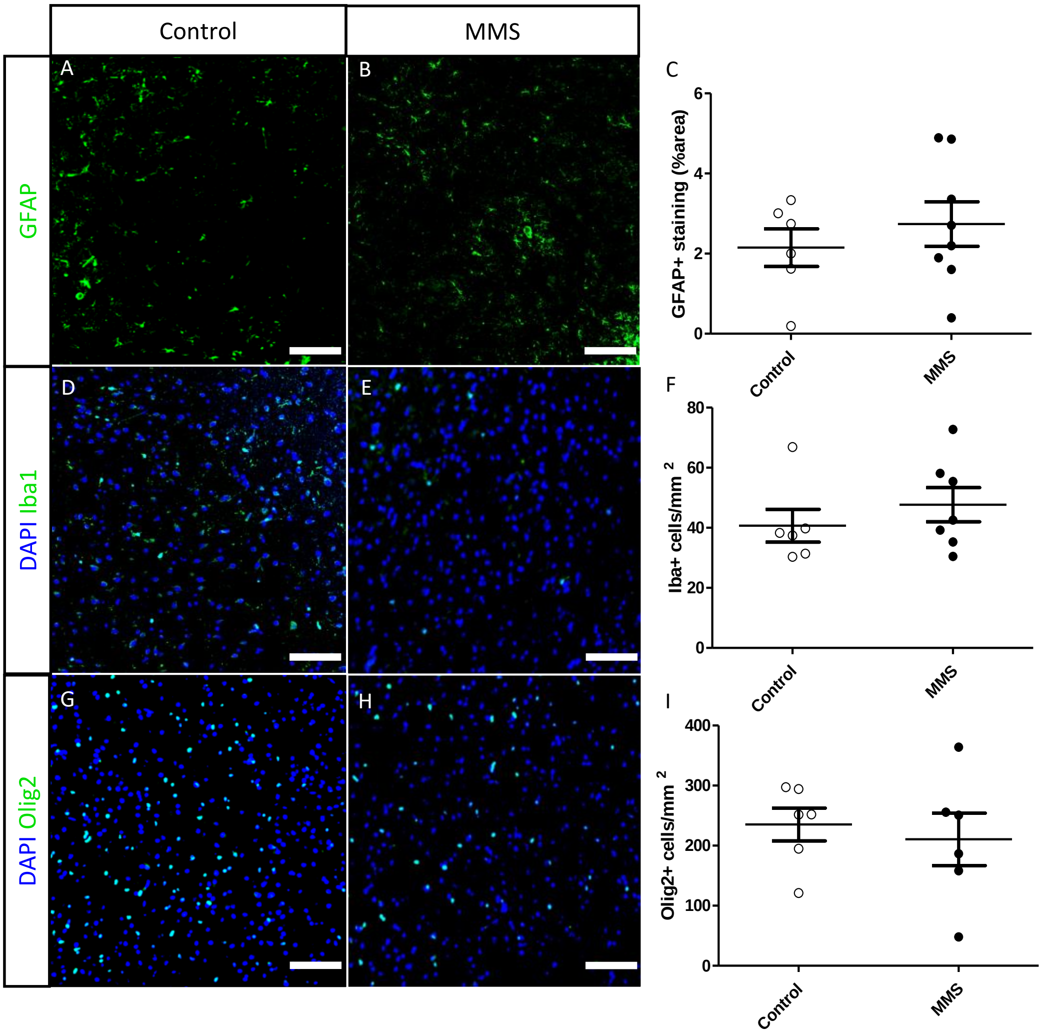

Supplementary Figure 2

A

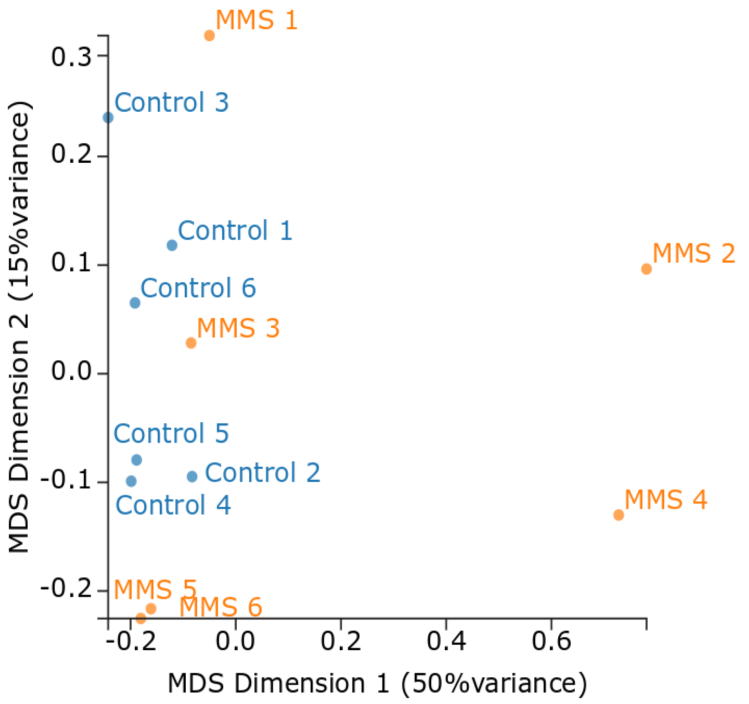


B

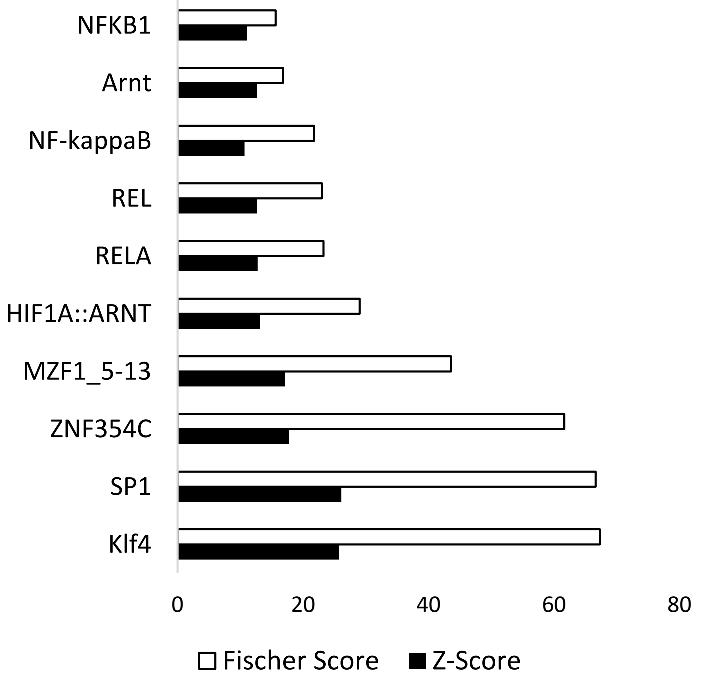

Supplementary Figure 3

A

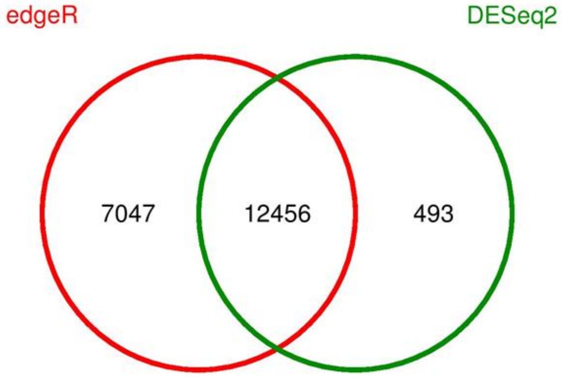


B

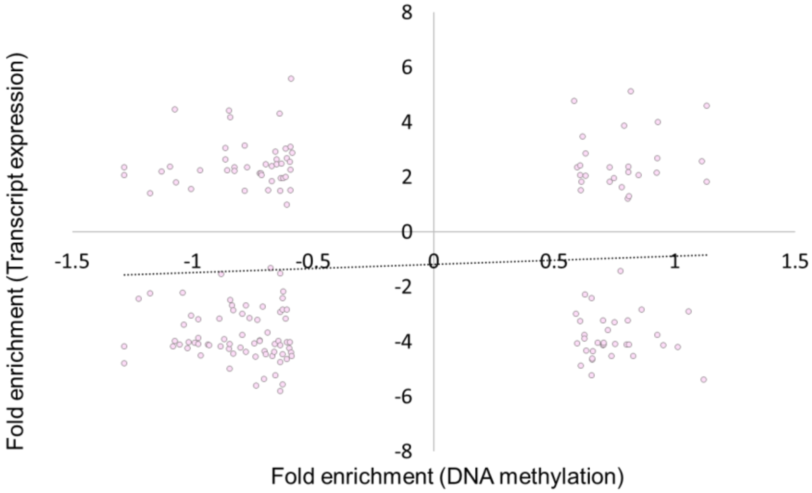

Supplementary Figure 4

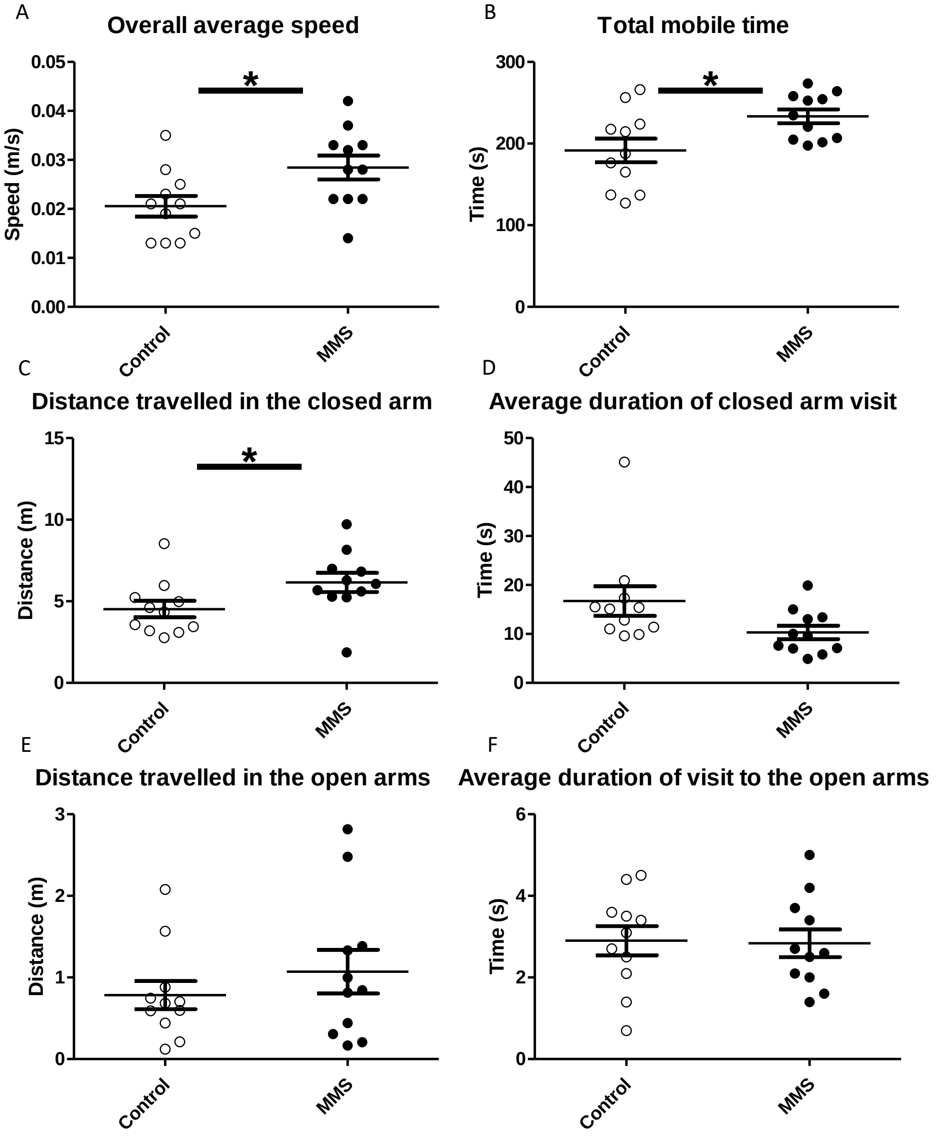

Supplementary Figure 5

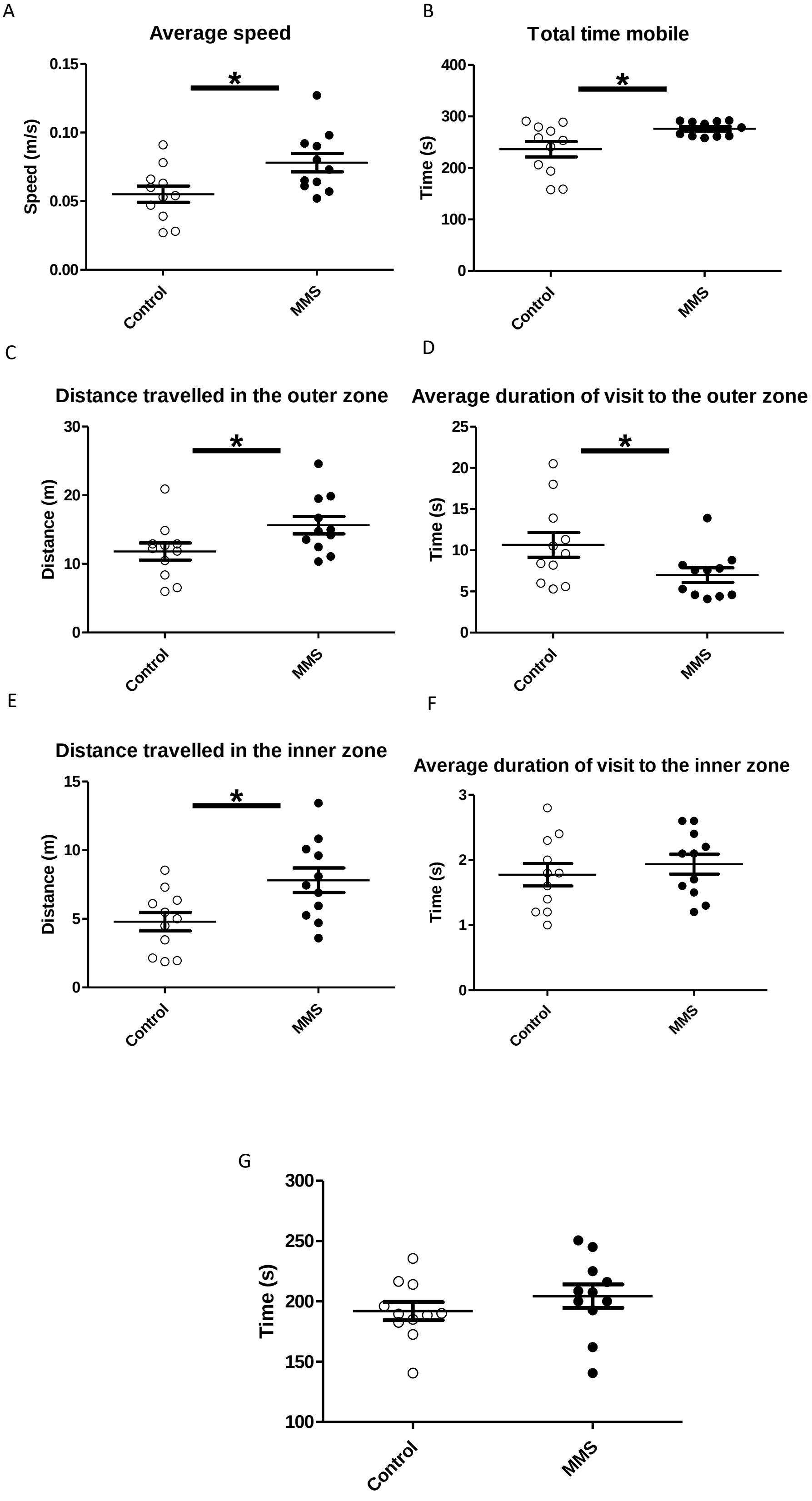

Supplementary Figure 6

A

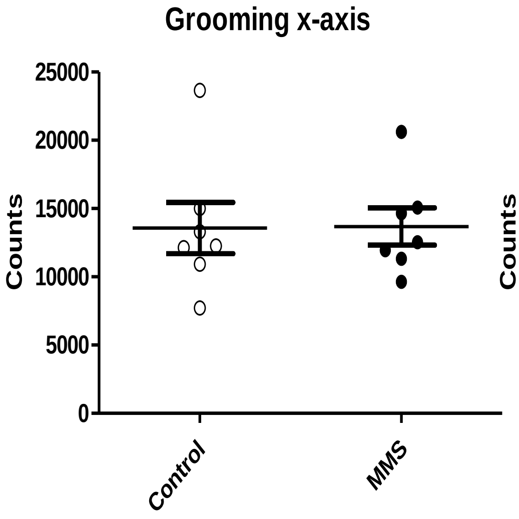


B

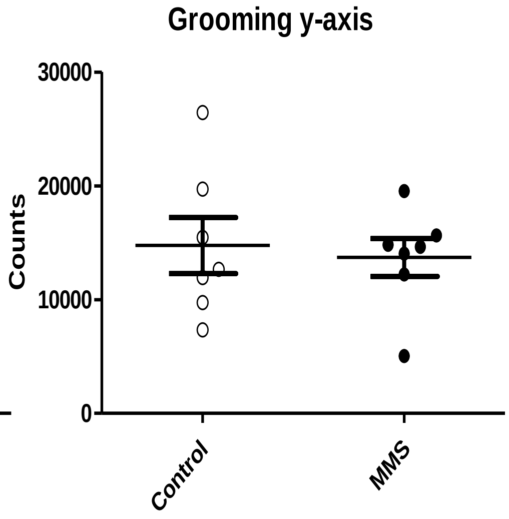


C

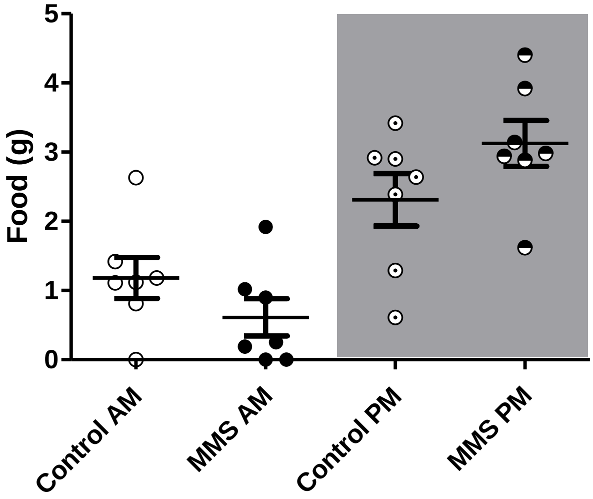


D

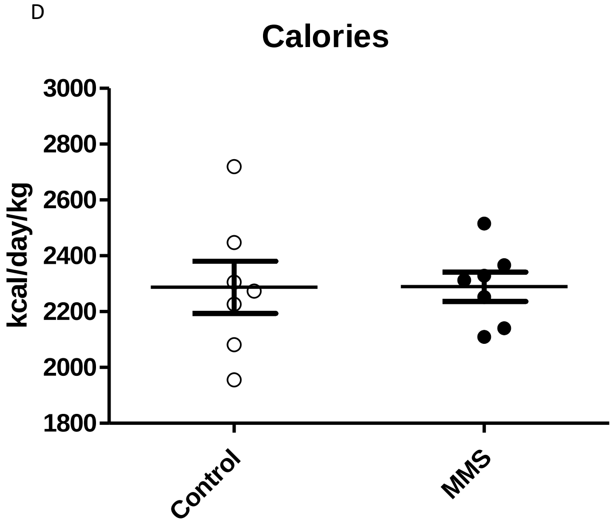


D

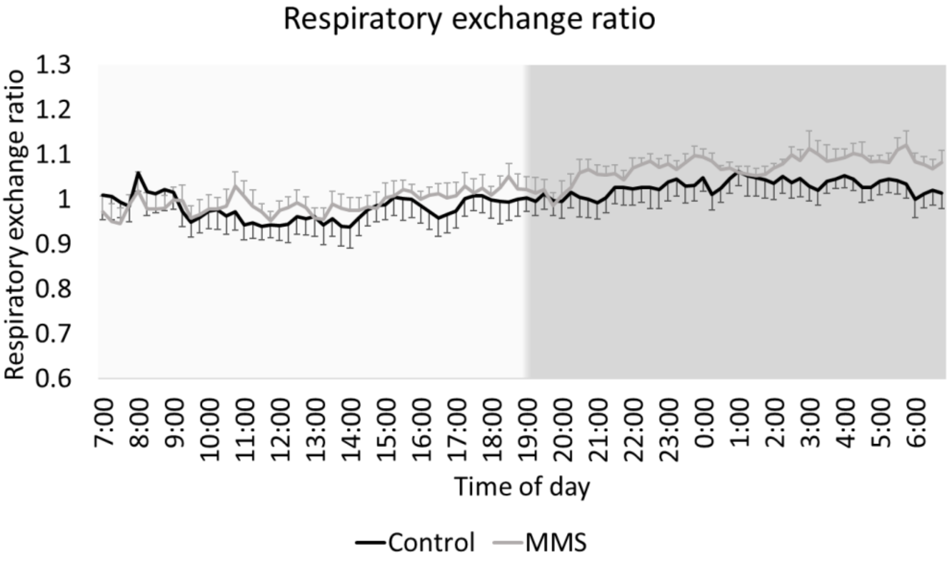

Supplementary Figure 7

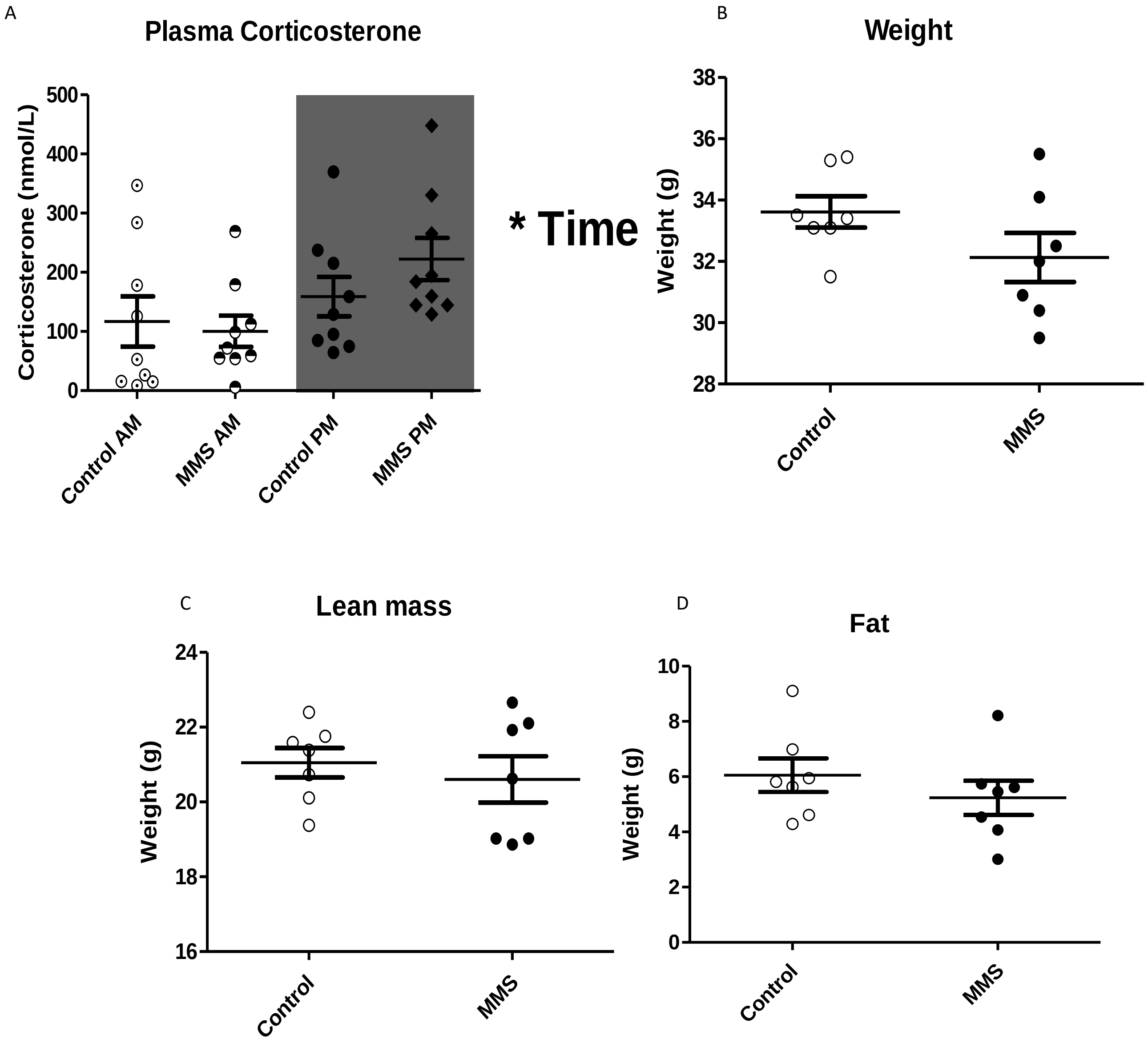

Supplementary Figure 8

A

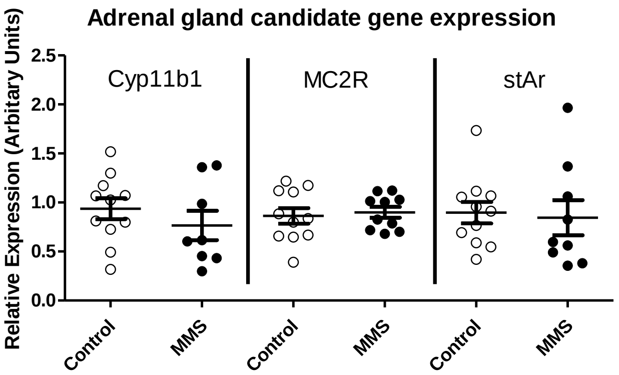


B

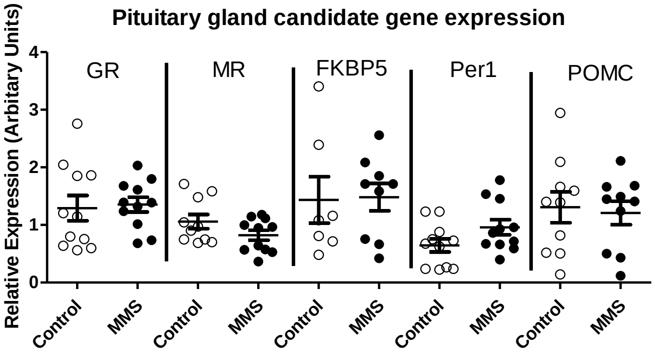


C

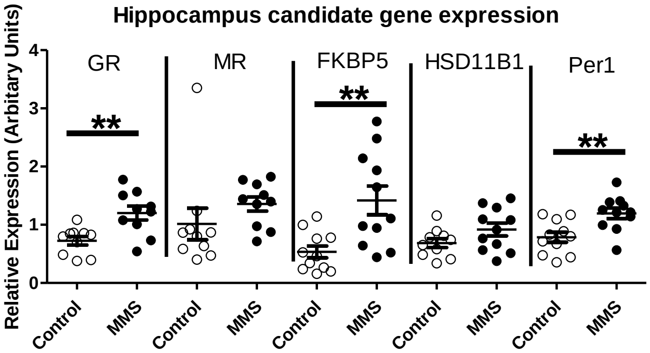
